# Hemodynamic forces can be accurately measured *in vivo* with optical tweezers

**DOI:** 10.1101/150367

**Authors:** Sébastien Harlepp, Fabrice Thalmann, Gautier Follain, Jacky G. Goetz

## Abstract

Force sensing and generation at the tissular and cellular scale is central to many biological events. There is a growing interest in modern cell biology for methods enabling force measurements *in vivo.* Optical trapping allows non-invasive probing of pico-Newton forces and thus emerged as a promising mean for assessing biomechanics *in vivo.* Nevertheless, the main obstacles rely in the accurate determination of the trap stiffness in heterogeneous living organisms, at any position where the trap is used. A proper calibration of the trap stiffness is thus required for performing accurate and reliable force measurements *in vivo.* Here, we introduce a method that overcomes these difficulties by accurately measuring hemodynamic profiles in order to calibrate the trap stiffness. Doing so, and using numerical methods to assess the accuracy of the experimental data, we measured flow profiles and drag forces imposed to trapped red blood cells of living zebrafish embryos. Using treatments enabling blood flow tuning, we demonstrated that such method is powerful in measuring hemodynamic forces *in vivo* with accuracy and confidence. Altogether, this study demonstrates the power of optical tweezing in measuring low range hemodynamic forces *in vivo* and offers an unprecedented tool in both cell and developmental biology.

## Introduction

Integration of biomechanics in cell biology studies has grown exponentially in the past years. The early observation that mechanical signals drive and modulate signaling pathways within cells through mechano-transduction has led to a paradigm shift when approaching cell biological phenomenon. Developmental biologists have led the way and undertook a series of mechanical manipulation and measurement to grasp the interplay between biomechanics and tissue morphogenesis (Petridou et al., 2017). Accurate manipulation, measurement and quantification of tissular and cellular forces *in vivo* are essential to such task. There are several research fields where quantification of forces *in vivo* is central. For example, the progression of many diseases such as cancer, cardiomyopathies, and myopathies is tightly linked to biomechanics. Much attention has been given to the interplay between static forces such as tissue elasticity and tumor progression (Jain et al., 2014). However, the biological ramifications of fluid forces, in development (Freund et al., 2012) and disease progression (Provenzano and Hingorani, 2013), are obvious and methods are needed for fine probing of these forces *in vivo*. Metastatic extravasation of circulating tumor cells (CTCs) is strongly influenced by mechanical inputs such as shear and adhesion forces, as well as vascular architecture (Azevedo et al., 2015; Wirtz et al., 2011). Similar to the extravasation of immune cells in an inflammatory context, efficient adhesion of CTCs to the vascular wall is needed before engaging extravasation (Reymond et al., 2013). While experiments have been largely performed *in vitro* (Chang et al., 2016; Labernadie et al., 2017), there is a growing need for establishing similar approaches *in vivo,* enabling high-accuracy measurements of forces in realistic physio-pathological situations. Performing such experiments *in vivo* faces however two major hurdles: it should be non-invasive and be able to reach deep tissues. Such limitations prevent the use of several contact techniques such as Atomic Force Microscopy (AFM), but led the development of non-invasive and non-contact tools such as magnetic tweezers (Brunet et al., 2013; Desprat et al., 2008) or optical tweezers (Ashkin and Dziedzic, 1987; Horner et al., 2017).

Optical tweezers suffer from the inconvenience of calibration *in vivo* and from the limitation in the application in deep tissues. While trapping cells in vivo, one needs to calibrate the set-up for every single cell that is trapped. Indeed, heterogeneity of cell size and of the refractive index prevents from applying a single calibration to multiple situation. Nevertheless, optical tweezers permit a dynamic analysis of the mechanical properties of cells (Huang et al., 2003; Monachino et al., 2017) and tissues (Gao et al., 2016; Lopez-Quesada et al., 2014) without altering embryonic development. Optical tweezing thus offers a wide palette for trapping and potentially measuring fluid forces *in vivo*. Recent pioneer work has shown that Red Blood Cell (RBC) can be trapped using optical tweezing in capillary vessels from mouse ears (Zhong et al., 2013). In this study, the authors moved the trapped RBC through the capillary vessel, and thereby induced artificial clots in the circulation. Although very powerful and promising, such technique remains limited when studying events deeply located in tissues.

The recent emergence of the zebrafish (ZF) embryo allows, due to its relative optical transparency, the use of a wide plethora of optical tools. Taking advantage of that property, Johansen et al. recently demonstrated the power of optical tweezing in a living ZF embryo (Johansen et al., 2016). They trapped and displaced multiple cell types in the blood flow, such as RBCs or macrophages, brought these cell types in contact with the vascular wall. Although this interesting study shows a large panel of potential applications, it does not provide an accurate quantification of the range of forces exerted or applied on the different objects. Few years before, we used a similar approach for quantifying the importance of the viscoelasticity of the arterial walls in ZF embryos by trapping RBCs at different positions of the vasculature (Anton et al., 2013). To our knowledge, this study was the first providing quantitative values from optical tweezing *in vivo*. We further used this approach to address the adhesion of epicardial cells to the pericardium during cardiac development in the zebrafish embryo (Peralta et al., 2013). More recently, oscillating optical tweezers deforming cell junctions on the drosophila embryo (Bambardekar et al., 2015; Sugimura et al., 2016) allowed the authors to accurately measure the tension forces between adjacent cells around 44pN.

Nevertheless, all these studies face a major obstacle, i.e. the calibration of the optical trap, which limits its use in realistic *in vivo* contexts. Recently, calibration of optical tweezers *in situ* has been achieved (Staunton et al., 2017). We provide here a mean to achieve accurate calibration in the context of probing hemodynamic forces in the zebrafish embryo. We first describe how to reach and quantify the physical parameters of the system by using high-speed imaging methods combined with image processing. Then, we describe how trapping of RBCs at different positions in the vasculature is processed in order to quantify the trap stiffness and to calculate the associated forces. We provide a numerical approach allowing us to solve the differential equation of the movement and compare the numerical data with the experimental ones to confirm the power of such calibration. Finally, we validate our approach by tuning the heart pacemaker activity and by measuring its impact on hemodynamic forces.

## Results and Discussion

### Fine measurement of velocity profiles for calibration of optical tweezers *in vivo*

Accurate calibration of optical tweezers *in vivo* requires a fine assessment of the physical parameters over time and space. Here, we introduce a method for accurate measurements of hemodynamic profiles in the zebrafish embryo (ZF). Hemodynamic profiles are measurements of velocity as a function of time, at a clearly located position in the ZF embryo. Such a method could be very useful in understanding the role played by hemodynamic forces in vascular morphogenesis, but also in pathological scenarios such as intravascular arrest and extravasation of circulating tumor cells (CTCs) or immune cells. We focused our analysis on the caudal plexus region of the ZF embryo (Fig.1A) whose optical characteristics and relative 3D structure make it optimal for highspeed imaging of the blood flow. We first perform high-speed imaging of the blood flow, at 200 frames per second (fps) and at intermediate magnification. Such a frame rate is optimal for fine tracking of individual RBCs in the vasculature (Movie S1). Raw acquisitions are processed to enhance the contrast of RBCs in the blood flow (Fig.1B, Movie S1). Subsequently, the processed data set is analyzed through a PIV (Particle Imaging Velocimetry) plugin developed on ImageJ^®^ and which is well-suited for automatic analysis of the blood flow movie (Fig.1B, Movie S1). The PIV allows a fine measurement of the velocity amplitudes and profiles at any position within the vasculature at a given time. Such a step is of utmost importance while studying hemodynamics with a Poiseuille-type distribution where velocity profiles vary significantly along a cross-section of a single vessel (Fig 1B). However, PIV analysis requires several parameters to be adjusted manually. While these parameters can potentially influence the measured velocities, we undertook a parallel approach aiming to extract the flow profiles of several RBCs using manual tracking performed in Fiji and rendered using Imaris software (Fig.1C, Movie S2). Because most of the RBCs follow the central streamline (Amini et al., 2014), manual tracking of individual RBCs allows to determine the flow profiles in the center of the measured vessel. We then compare these profiles to the ones obtained upon PIV treatment. The comparison of both methods allows us to estimate the accuracy of our measurements, which is around 50μm/s for the determination of the blood flow velocity. We then fit the PIV with theoretical profiles (Fig.1C). The function used is adapted from our previous work (Anton et al., 2013) and can be written as follows:

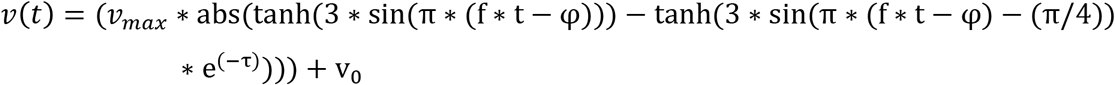

v_max_ represents the maximal velocity, f the heart beating frequency, φ the phase shift representing the arbitrary origin of the experimental signal, v_0_ the minimal velocity and τ the damping during the diastolic phase linked to the endothelial barrier viscoelastic behavior.

**Figure 1:**
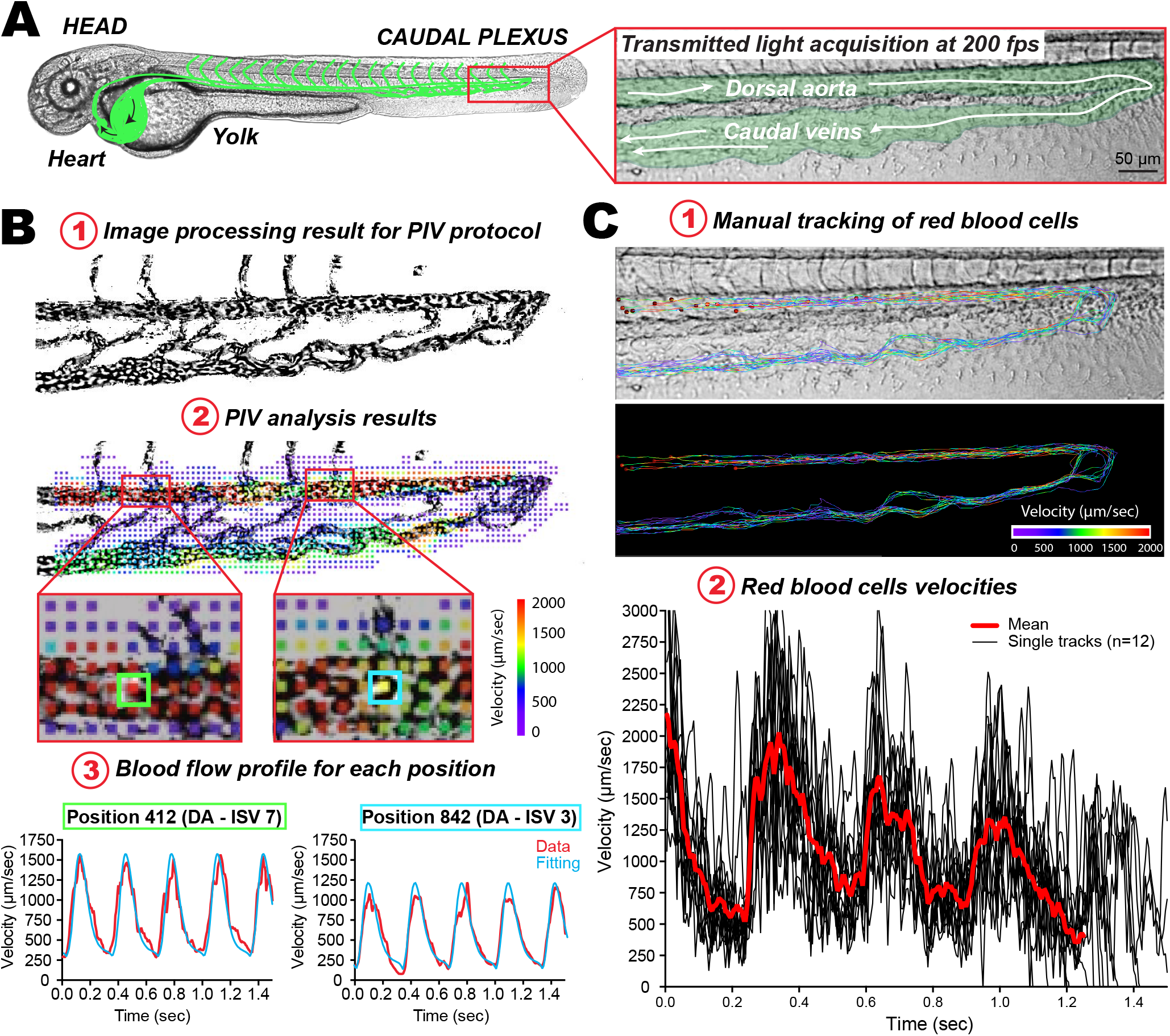
Fine measurements of blood flow velocities in the zebrafish embryo. **A.** Experimental workflow: a representative image of a 48hpf zebrafish embryo image is presented (the vasculature network is schematized in green) and a zoom of the caudal plexus region is presented in a separated box. **B.** PIV analysis: 1. the images resulting from high-speed acquisition of the blood flow (200 fps) were processed such that high contrast is obtained for circulating RBCs. 2. The PIV analysis provides a color-coded velocity map over the entire image, in any region of interest. Flow profiles can thus be extracted at any given position. This further allows theoretical analysis of the flow values, which will be used in the simulation used for measuring the trap stiffness. 3. Here, two positions, with distinct flow profiles and separated by roughly 250 μm, are presented. Note the different flow profiles that can be observed and fitted. **C.** Particle tracking analysis. Manual tracking of individual RBCs in perfused vessels can be performed 1. Several tracks obtained over 12 RBCs are displayed. Tracks are color-coded according to their instantaneous velocity. Note the higher values and pulsatility obtained in the dorsal aorta. 2. Instantaneous velocity frame after frame is plotted over a track spanning the entire caudal plexus, for 12 RBCs.

Combining these different methods allow a very accurate assessment of velocity profiles at any position within the vasculature. This further permits to model the profile theoretically and to adjust an equation whose parameters can be further used in the numerical simulation. Nevertheless, before performing numerical simulation of blood flow behavior in the ZF embryo, one should assess the efficacy of optical tweezing of circulating RBCs in developing blood vessels.

### Optical trapping of RBCs, image analysis and photodiode measurements

Although it is technically possible to trap RBCs (or other cells) anywhere in the zebrafish embryo (see Peralta et al., 2013), we mostly performed optical tweezing in the caudal plexus. Here, in comparison to high-speed recording of the blood flow, a high magnification and numerical aperture objective is used to optically trap the RBCs in the flow (Fig.2A, Movie S3). These trapped RBCs are mostly subjected to the dragging force of the blood flow and the restoring force of the optical tweezers (Jun et al., 2014). We record the displacement of the RBCs within the optical trap on a quadrant photodiode at high frequency (over 4 kHz) (Fig.2B). This temporal acquisition mirrors the cellular displacement from the trap center and is expressed in Volt. Here, we convert this displacement expressed in volts to displacement expressed in μm by applying a conversion factor extracted from the 2 curves, allowing to superimpose this curve to the displacement measured with the camera. We further compute it as the power spectral density of the fluctuations (Fig.2B - black track), over which, one can fit the optical tweezers behavior of a trapped cell in vitro in absence of flow (supplementary information). The parameters that are obtained allow us to extract a first approximation of the trap stiffness which is k=4.7*10^-5^ N/m. A similar temporal trapped RBC acquisition is simultaneously performed at 200 fps. The resulting kymograph shows the cellular displacement within the trap (Fig.2C) where accurate distances can be extracted (μm). From the cellular displacement in the trap and the measured stiffness, one can now convert the displacement track into an accurate force measurement (Fig.2D). When repeating the acquisition over the caudal plexus of the 2dpf ZF embryo, we are now capable of drawing a hemodynamic force map (Follain et al., submitted, personal communication). From the forces extracted with this method, it is also possible to elaborate the flow profile and thus to extract the shear stress one cell has to overcome once attached to the endothelium. In addition, one can now observe and highlight the Poiseuille flow profile of the developing dorsal aorta of a 2 dpf ZF embryo (Fig.2E). This allows quantifying both the velocity and the shear stress in close proximity to the endothelial walls. It is important to note at this stage that the calibration of optical tweezers was here performed in a blood vessel with pulsatile flow (dorsal aorta). Such a parameter could greatly influence the measured trap stiffness, as well as the ways to measure it. However, our workflow now allows conducting numerical simulations that can be compared to the previous data.

**Figure 2:**
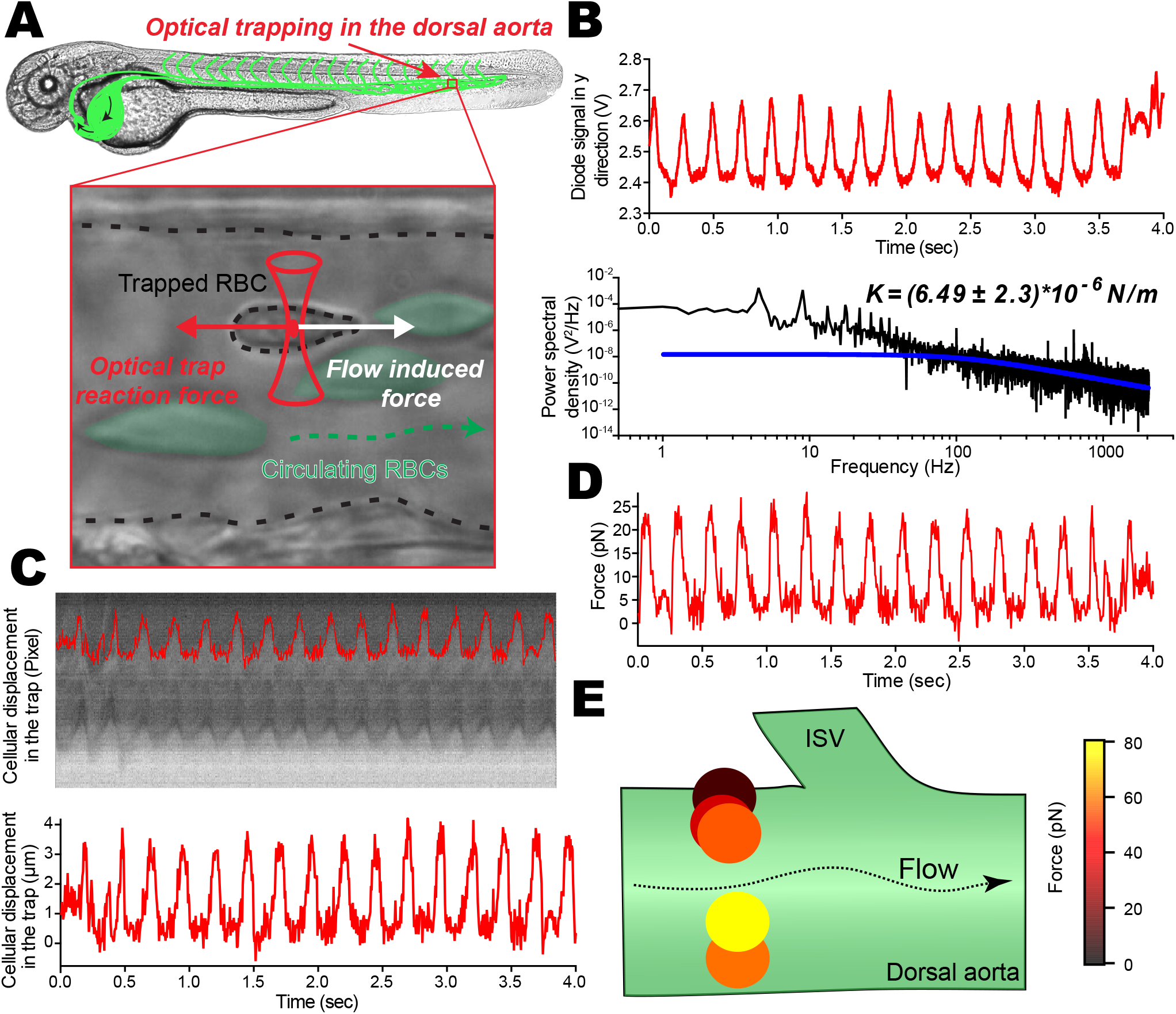
Optical trapping of circulating RBCs, image analysis and photodiode measurements. **A.** Scheme of the experiment where the optical trap is located in the caudal plexus (dorsal aorta) of the zebrafish embryo. The trapped RBC is subjected to the blood flow-driven dragging force as well as to the restoring force from the optical trap. **B.** Thanks to the flow pulsatility occurring in arterial vessels, the time trace of the optically trapped RBC is followed on the quadrant diode. The time trace is further treated to access the power spectrum (black curve), revealing peaks of pulsatility as well as a corner frequency f_c_ around 250 Hz. This is highlighted with the theoretical curve for steady optical trapping experiments (blue curve). The cut off frequency, obtained from a Lorentzian approximation, provides a first approximation of the trap stiffness, which is about 4.7*10^-5^N/m **C.** A kymographic analysis of a single trapped RBC imaged at 200 fps is performed. Kymographs allow fine tracking of RBC displacement within the optical trap, thereby providing a real time track in μm. **D.** A mirror force track over time is obtained and results from the use of the theoretical trap stiffness derived from the cut off frequency (theoretical curve). **E.** Any given position in the zebrafish vasculature can be probed, providing an accurate measure of hemodynamic forces. Here, a few trapping RBCs in a cross-section of the DA allow to appreciate the behavior of a Poiseuille-type of flow.

### Performing numerical simulations for accurate measurement of the trap stiffness

Before starting running simulations of *in vivo* experiments, one needs to verify and assess the robustness of the code (see supplementary methods). We first performed simulations in absence of external flow and tuned the trap stiffness from 10^-5^ to 10^-4^ N/m (Fig.3A). The model predicts that an increase in the trap stiffness should lead to a decrease in the fluctuations of the cell within the trap as well as an increase in the cut off frequency. We next considered the presence of a pulsatile flow for which we imposed a fixed velocity, frequency and damping values while only tuning the trap stiffness again from 10^-5^ to 10^-4^ N/m, The flow parameters are the following: maximal velocity of 1300 μm/sec, frequency of 2.1Hz and the damping τ=0.8 (this latter value correspond to an average value obtained through the caudal plexus) (Fig.3B,C). As expected, the trap stiffness affects the cut off frequency in a similar manner to the ones obtained in absence of flow (Fig.3B). In addition, we observed and measured a significant decrease in the peaks amplitude with increasing trap stiffness confirming that the fluctuations of the cell within the trap are directly linked to the stiffness (Fig.3B). The simulation allows us to extract also the displacement of the RBCs over time and these displacements clearly demonstrated the impact of the trap stiffness on the cellular movement (Fig.3C). More precisely, we observed a displacement of 7.7μm within the trap when the stiffness was set to 10^-5^N/m, and a displacement of 770nm when the stiffness was set to 10^-4^N/m. A cellular displacement of 7.7μm within the trap, which exceeds the radius of individual RBCs, will inevitably lead to the RBC escaping from the trap. Such simulations confirm that stiffness of the trap impacts the cut off frequency and linearly impacts the overall amplitude of fluctuations at low frequencies that are mainly driven by the amplitude of the flow. It is important to note at this stage that these simulations were run by averaging the spectral density over a hundred runs. A major bottleneck of such an experiment is however the acquisition time, which should be as low as possible to exclude or prevent external perturbations such as drift of the sample/embryo or trapped RBCs that are being ejected from the trap by collision from other circulating RBCs. We therefore assessed the sensitivity of this method by modulating the number of temporal averages from 100 to 1 while keeping all the other parameters constant. We ran a first series of simulations at high trap stiffness (k=10^-4^N/m) and observed that the trap cut off frequency remains well defined when compared to the flow-driven frequency dependence (Fig.3D). We thus concluded that the number of acquisitions is irrelevant. Nevertheless, in a second experiment where the trap stiffness was set to 5*10^-5^N/m (Fig.3E), we observed that such stiffness leads to a trap cut off frequency that cannot be extracted from the flow-driven frequency dependence. Here, a simulation averaging of 100 times is required to reach the trap cut off frequency. In other words, probing forces at such parameters is incompatible with experimental recording that would require acquisition of about 100 seconds or more. For this reason, we rely now on the developed numerical code (supplementary material) as well as on fine flow mapping described previously (Fig.1). A combination of numerical code and fine flow mapping allows us to calibrate the optical trap from the experimental data by applying a fit to the obtained curves. First, we extract the experimental data from the signal obtained on the quadrant diode and transform them into a real displacement in μm (Fig.3F, red plot). We run the simulation with the flow parameters extracted from the previous PIV experiment and adjust the trap stiffness until the simulated displacement adjusts the experimental one (Fig.3F, blue plot). This theoretical curve gives a good approximation on the beating frequency and the endothelial damping. The values obtained from fine flow mapping (PIV + tracking, Fig.1) are fed in the numerical simulation as input parameters before running the fit and superimpose the power spectral density curves obtained experimentally (Fig.3G, red) and theoretically (Fig.3G, black). This example demonstrates the good accordance between velocity peaks induced by flow pulsatility and the appearance of the cut off frequency around 250Hz. Altogether, this demonstrates that the calibration of the set-up is achieved. This gives now access to fine blood flow force mapping at any position in the developing ZF embryo.

**Figure 3:**
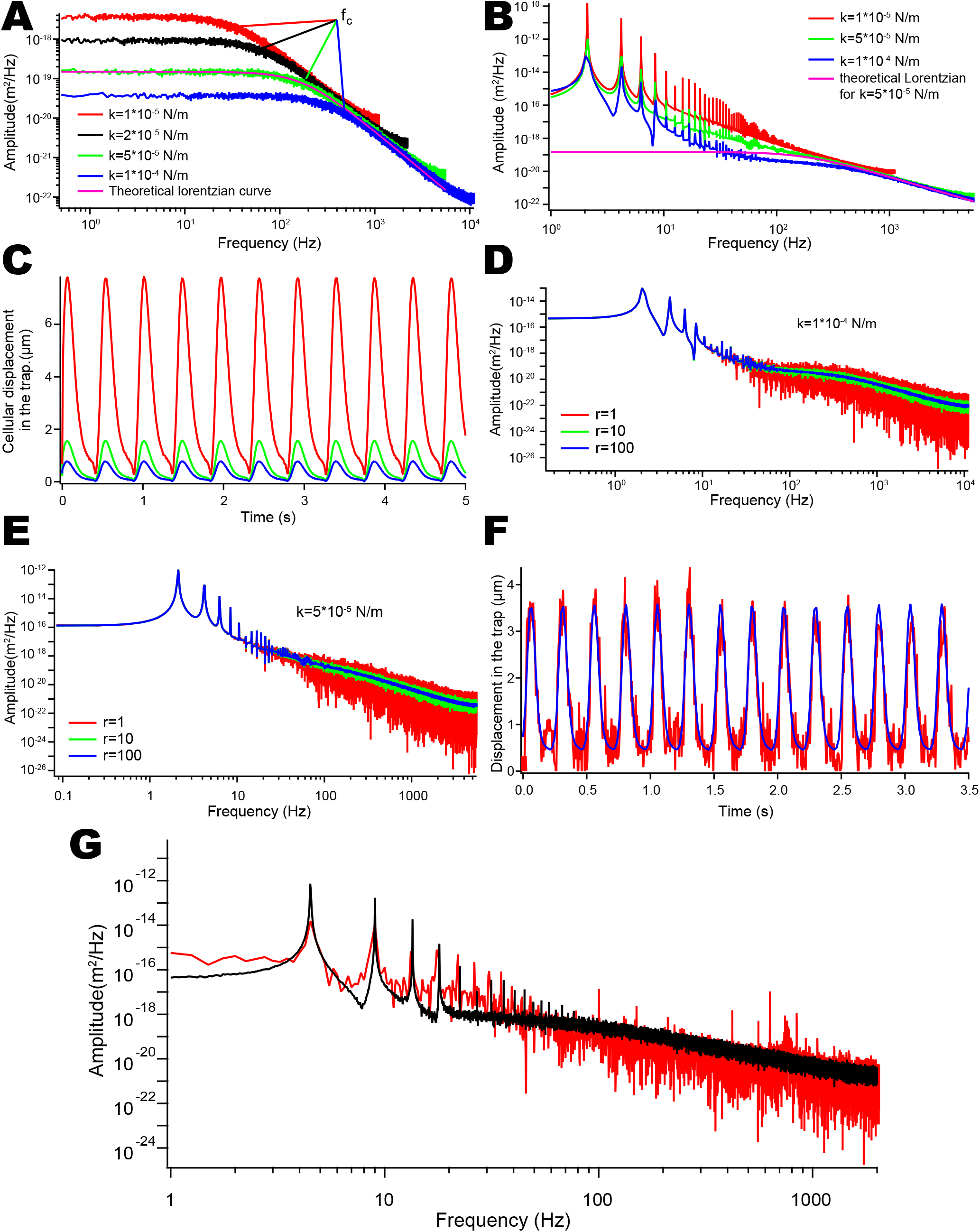
Numerical simulations. **A.** Power spectrum of different tracks is represented graphically. No external pulsatile flow was applied and the trap stiffness was modulated from 10^-5^ to 10^-4^ N/m to show that global fluctuations in the trap decrease when the stiffness was increased (ordinate at the origin) and that the cutoff frequency increases to higher frequency from 50 to 350 Hz. **B.** Power spectral representations of different tracks resulting from same pulsatile flow. The trap stiffness is modulated from 10^-5^ to 10^-4^ N/m. The cutoff frequency increases to higher frequency when the stiffness increases as shown in A. The amplitude of the peaks at 2 Hz and its harmonics diminishes when the stiffness increases. **C.** Movement of RBCs within the optical trap is tracked and plotted over time. Low trap stiffness allows trapped RBCs to reach 6μm oscillations that are bigger than the cellular diameter and would lead to RBCs escaping the trap over time. **D.** Three different power spectrum with constant velocity and physical parameters k=5.10^-5^N/m, but with different spectral averaging from 1 to 100, are represented. While the signal to noise ratio is increasing as the square of the averages the cutoff frequency due to the trap stiffness was visible without averaging. **E.** Three different power spectrum with constant velocity and physical parameters k=5.10N/m, but with different averaging from 1 to 100. As in D, the signal to noise ratio is increasing as the square of the number of averages nevertheless the cut off frequency (around 150Hz) is hardly visible even after 100 spectral averages. This is due to the overlap of the flow contribution in the power spectrum and the cutoff frequency due to trap stiffness making its distinction complicated. **F.** Displacement of a single RBC within the optical trap is tracked and plotted over time. The blue curve represents the theoretical function of the flow pulses, which perfectly matches the experimental data. The obtained function is later used to model and fit the experimental power spectrum. **G.** Experimental power spectrum is obtained from a trapped RBC. The black curve represents the fitting of the experimental red curve with the numerical model. Such fitting permits to extract with high accuracy the trap stiffness

### Blood flow tuning can be accurately assessed with OT *in vivo*

A proper calibration of the OT allows envisioning fine flow mapping in the ZF embryo. In order to demonstrate the power of such a method, we tuned the pacemaker activity using a previously described pharmacological treatment and measured its impact on flow profiles and applied forces. Lidocaine, a sodium channel blocker, has been shown to reduce significantly the pacemaker activity of 2dpf ZF embryos (Cummins, 2007; Vermot et al., 2009). Using high-speed imaging of heart beatings, we observed that Lidocaine reduced the pacemaker activity by 30-35% (data not shown, Follain et al., submitted, personal communication) and impacted the flow velocity accordingly. These behaviors are in good agreement with previous works performed with lidocaine (Anton et al., 2013; Heckel et al., 2015; Vermot et al., 2009). Optical tweezing of single RBCs was performed at two different positions in the caudal region of the dorsal aorta of control and lidocaine-treated embryos (Fig.4A-B). Prior to optical tweezing, flow profiles at each respective position were probed using high-speed imaging (200 fps) followed by PIV analysis. Flow mapping indicates the significant decrease in flow velocities imposed by lidocaine and the underlying reduction in pacemaker activity. Both flow pulsatility as well as velocity amplitude are reduced at both positions (Fig4.C-D). We then trapped single RBCs at the two positions and tracked the cellular movement within the trap over time (Fig.4E-F). Kymographic analysis revealed RBCs oscillations within the trap, which can be plotted over time (Fig 4G-H). The profiles that are obtained perfectly mirror the velocity profiles from the PIV analysis, with a marked decrease in both pulse frequency and flow amplitude. Nevertheless, as shown previously, the trap stiffness controls the amplitude of movement of RBCs within the trap. We thus adjusted the laser power such that RBCs would be hold in the trap without escaping in the control experiments. We thus increased the laser power in the control condition compared to the lidocaine condition. We then extracted the trap stiffness from the power spectra adjusted using our previously described numerical model and plotted the force tracks that are associated to cellular displacements (Fig.4I-J). Such an analysis allows to accurately measure flow forces and, in that case, to probe the impact of a reduction in pacemaker activity on hemodynamic forces at high accuracy.

**Figure 4:**
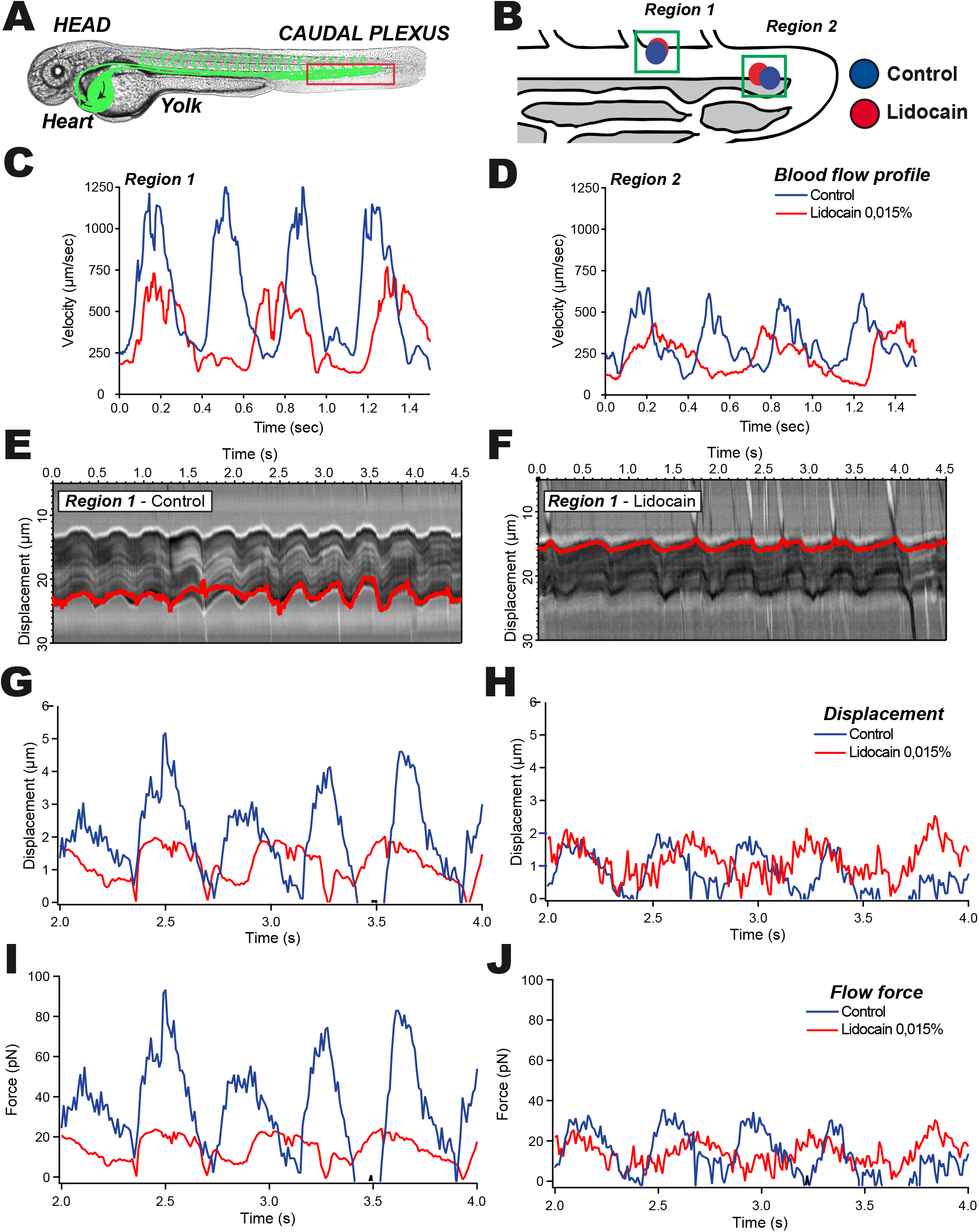
The impact of blood flow tuning on hemodynamic forces can be accurately measured using calibrated optical tweezing. **A.** A representative image of the zebrafish embryo at 48hpf is provided and the region of interest is highlighted in red. Blood flow tuning is achieved by pre-treating the embryos with lidocain (0,0015%). **B.** Two positions within the caudal region of the dorsal aorta are probed with optical tweezers. **C-D.** PIV tracks are obtained from high-speed imaging and analysis of region 1 (C) and region 2 (D). **E-F.** Kymographic analysis of the optically-trapped RBC in region 1 in control (E) and lidocain-treated embryos (F). The displacement track is represented in red. **G-H.** Displacement of the trapped RBC is measured and plotted in region 1 (G) and region 2 (H). **I-J.** Force curves from region 1 (I) and 2 (J) are extracted from the displacement of RBCs within the calibrated optical trap.

In conclusion, we have described here a fast and straightforward method to probe and measure, hemodynamic forces in vivo. By combining flow profile dissection, power spectrum acquisition and numerical analysis of the experimental data, we could extract the stiffness of the optical trap that is essential for correlating the trapping effect to the imposed force. The power of this method and analysis is that the numerical simulation provides a fast and reliable mean to assess trap stiffness, and the underlying forces. As described previously (Zhong et al., 2013) and detailed in the supplementary methods (Supplementary information), one can neglect inertia. Indeed, the temporal resolution of imaging of the blood flow is limited and leads to two major forces: the restoring force exerted by the optical trap to the trapped object (i.e a RBC) and the dragging force from the blood flow. These two forces have separated frequency responses. While the restoring force acts at high frequency (100-1000Hz), the dragging force, in the case of the ZF embryo, is closely linked the heart beat frequency (around 2-3Hz). In these conditions, previous work demonstrated analytically that two signals with frequency domains sufficiently separated could be solved in the Fourier domain as a sum of the two contributions (Seth Roberts et al., 2007). In their work they imposed oscillations at frequency far from the cut off frequency to the trapped bead with an electrical field. From the collected oscillation, they plotted the power spectrum presenting a defined peak at the oscillation frequency. The amplitude of this peak due to its distance from the cut off frequency was directly proportional to the number of charges created.

Our method suffers from two limitations. First, we consider RBCs as spherical objects from which the corresponding dragging force is derived. This consideration does not take into account the RBC deformations. Second, we do not precisely measure the distance separating RBCs from the vessel wall and thus consider a constant viscosity instead of considering Faxen’s law (Jun et al., 2014). We therefore introduce underestimations in the resulting calibration. These underestimations represent an error on the force of about 20% close to the endothelial wall.

Altogether, we have shown here that precise probing of hemodynamic forces using optical tweezing *in vivo* can be achieved. Such measurements, when applied to the understanding of hemodynamics in vascular morphogenesis (Sugden et al., 2017), atherosclerosis (Namdee et al., 2014), or thrombosis (Karachaliou et al., 2015) events could provide unexpected insights in the contribution of fluid mechanics. When using the trapping function of optical tweezers, one can also redirect circulation of RBCs in the ZF embryo and unravel the importance of vascular architecture in driving its perfusion (Sugden et al., 2017). Another important question that could be solved using this tool is the potential contribution of hemodynamics to intravascular arrest of immune or circulating tumor cells. Probing the shear stress in very close proximity to the vessel wall could help understanding whether rolling of immune or tumor cells, usually attributed to selectin adhesion (Brunk and Hammer, 1997; Lawrence et al., 1997; Sun et al., 2015), can be tuned by hemodynamics. During their journey in the blood circulation, CTCs are subjected to shear forces and collisions with host cells. Only CTCs that overcome or exploit these forces will eventually arrest in a vascular regions, preceding metastatic outgrowth. A major advantage provided by calibrated optical tweezing relies in the ability to probe hemodynamic forces precisely at the vessel wall. These regions cannot be reached with other classical flow mapping methods, although they are of utmost importance in the context of intravascular arrest of tumor or immune cells. Assessing, using calibrated optical tweezing, the impact of flow forces on such events, and many others, is thus within reach and could lead to very important observations in a near future.

## Acknowledgements

We thank all members of the Goetz Lab for helpful discussions. We thank Julien Vermot for the original experiments in the zebrafish embryo. We are grateful to Pascal Kessler (IGBMC) for his help with the Imaris software. This work has been funded by labex NIE (to S.H. and J. G.), Plan Cancer (OptoMetaTrap, to J.G. and S.H) and CNRS IMAG’IN (to S.H., J.G.) and by institutional funds from INSERM and University of Strasbourg. G.F. is supported by a Ph.D. fellowship from La Ligue Contre le Cancer.

## Supplementary Material

**Supplementary Methods:** We provide as a stand-alone supplementary method a detailed description of the numerical resolution of the Brownian equation.

**Movie S1: PIV analysis for assessing hemodynamic profiles.** Transmitted light highspeed acquisition (200 fps) of the blood flow in the caudal plexus of a zebrafish embryo (top panel). Acquisitions are optimized for PIV analysis (middle panel) and final PIV rendering (bottom panel).

**Movie S2: Manual tracking (and rendering) of RBCs for assessing the blood flow profiles.** Transmitted light high-speed acquisition (200 fps) of the blood flow in the caudal plexus of a zebrafish embryo (top panel). 12 RBCs are manually tracked all along the caudal plexus, tracks (and related velocities) are revealed (bottom panel).

**Movie S3: Optical tweezing of RBCs *in vivo*.** Optical tweezing of single RBCs in the dorsal aorta of a control (left panel) and a lidocain-treated (right panel) zebrafish embryo.

## Material and Methods

### ZF handling and mounting for high-speed imaging

*Tg(fli1a:eGFP)* Zebrafish (*Danio rerio*) embryos used in this study come from a Tübingen background. Embryos were maintained at 28° in Danieau 0.3X medium (17,4 mM NaCl, 0,2 mM KCl, 0,1 mM MgSO_4_, 0,2 mM Ca(NO_3_)_2_) buffered with HEPES 0,15 mM (pH = 7.6), supplemented with 200 μM of 1-Phenyl-2-thiourea (Sigma-Aldrich) after 24hpf. At 48hpf, embryos were freed from their chorion and mount in an agarose drop (0,8%) deposite on a glass-bottomed dish (MatTek) compatible with high-resolution imaging. The embryos were positioned in the microscopic framework in order to overlap the DA longitudinal axis with the *x-axis* and the ISV with the *y-axis*. For lidocaine treatment experiments, embryos were incubated in Danieau with 0,0015% lidocaine vs control vehicle (EtOH) for 2 hours before mounting and imaging. Pacemaker activity of the heart is assessed using ane USB 3.0 uEye IDS CCD camera (IDEX) mounted on a DMIRE2 inverted microscope (Leica) using transmitted light. Heartbeats were acquired at 80 frames per second (fps). Kymographic analysis was performed for extracting the beating frequency.

### High-speed acquisition and Particle Image Velocimetry (PIV) analysis

For the PIV analysis, high-speed recordings of the blood flow were performed using a Thorlabs DCC3240C Cmos camera at 200 fps mounted on the inverted microscope coupled with a UPLFLN 20X/0.4 objective (Olympus). The red blood cell positions at each frame were cross-correlated to the following frame using an adapted Particle Image Velocimetry (PIV) protocol from ImageJ software available online (https://sites.google.com/site/qingzongtseng/piv). The individual displacements as well as the associated frame rate give then access to the individual velocity profiles of the blood cells in the bloodstream. These results were compared with results generated from Manual particle tracking (ImageJ) sequences on the same movies. This allows to finely tune the PIV parameters in order to access to the velocity distribution in the vasculature.

### Optical trapping

A 1064 nm laser (Navigator 1064-7Y Spectra Physics) feeds the back aperture of the UPLFLNP 100X/1.3 objective (Olympus) to generate optical trap, the objective is mounted on a thermostated inverted microscope. Trapping of red blood cells was performed as described in Anton (Anton et al., 2013) in a thermoregulated chamber ensuring the embryos to remain at 28°C. We acquired the displacement with the Thorlabs Cmos camera at 200 fps and simultaneously acquire the temporal signal on a quadrant photodiode at 20kHz to calibrate the setup to reconstruct the power spectrum and thus allowing the calibration of the trap.

### Numerical simulation

A full and detailed description is provided in the supplemental material (see supplementary methods). A general Langevin equation describing the Brownian evolution of a particle combined to an external flow reads

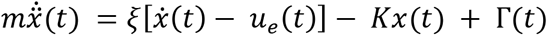

In the present case, let us assume that the parameters are ξ = 4,5*10^-8^ N.s.m^-1^ the Stokes friction coefficient of the cell, K = 1*10^-4^ N.m^-1^ the optical tweezer spring stiffness, u_e_(t) the external pulsatile flow with a velocity of the order of a few mm.s^-1^, and m = 2.1*10^-14^ kg the mass of a blood cell of radius 1.5 μm and volume mass 1500 kg/m^3^ (in absence of hydrodynamic corrections). T(t) is the random noise, or Langevin force. This random force is supposed to be white, stationary and Gaussian, with the standard time correlations.

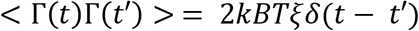

The overall process is Markovian. Considering now the effect of periodic external force acting on the cell, one observes that three different regimes emerge, delimited by two characteristic frequencies ω_s_/(2π) and ω_i_/(2π). For ξω_s_ = K, the friction force matches the tweezer restoring force, while for ξω_i_ = mω_*i*^2^_ friction and inertia come even. Straightforward numerical estimates lead to ω_s_/(2π) = 350 Hz and ω_i_/(2π) = 340 kHz.

At low frequencies (f < 300 Hz), the position of the cell in the optical trap follows closely the external drag force ξ.u_e_. At these frequencies, the trapped cell follows the regime dominated by the external flow and the optical trap stiffness. Inertia can be safely neglected in the range of frequencies f < 10^4^ Hz for which experimental sampling is done. At frequencies between 300 and 10^4^ Hz, the position of the cell in the optical trap is dominated by thermal fluctuations and is represented on the power spectrum by a −2 power law. The frequency where the cellular position switches from the dragging and trapping regime to the thermal fluctuations regime is called the cut off frequency.

We therefore consider the overdamped, or Smoluchowsky, limit of the Langevin equation

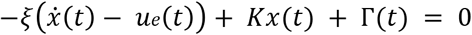

This equation represents the master equation used to run the numerical simulations.

